# Preterm birth alters the maturation of the GABAergic system in the human prefrontal cortex

**DOI:** 10.1101/2021.12.14.472624

**Authors:** Helene Lacaille, Claire-Marie Vacher, Anna A. Penn

**Affiliations:** Department of Pediatrics, Columbia University, New York-Presbyterian Morgan Stanley Children’s Hospital, New York, NY, USA

**Keywords:** Prematurity, GABA, prefrontal cortex, neurodevelopment, maturation, astrocyte

## Abstract

Developmental changes in GABAergic and glutamatergic systems during frontal lobe development have been hypothesized to play a key role in neurodevelopmental disorders seen in children born very preterm or low birth weight, but the associated cellular changes have not yet been identified. Here we studied the molecular development of the GABAergic system specifically in the dorsolateral prefrontal cortex, a region that that has been implicated in neurodevelopmental and psychiatric disorders. The maturation state of the GABAergic system in this region was assessed in human post-mortem brain samples, from term infants ranging in age from 0 to 8 months (n=17 male, 9 female). Gene expression was measured for 47 GABAergic genes and used to calculate a maturation index. This maturation index was significantly more dynamic in male than female infants. To evaluate the impact of premature birth on the GABAergic system development, samples from one-month-old term (n=9 male, 4 female) and one-month corrected-age (n=8 male, 6 female) very preterm infants, were compared using the same gene list and methodology. The maturation index for the GABAergic system was significantly lower (–50% p<0.05) in male preterm infants, with major alterations in genes linked to GABAergic function in astrocytes, suggesting astrocytic GABAergic developmental changes as a new cellular mechanism underlying preterm brain injury.

## Introduction

Premature birth, affecting 10% of live births, is a leading cause of neonatal morbidity that can induce long-term neurological impairment (Adams-Chapman & Stoll, 2006; Arpino et al., 2010; Blencowe et al., 2013; Knuesel et al., 2014; Marín, 2012; Meldrum et al., 2013). Extensive alterations in frontal lobe development have been reported in children born very preterm (<32 weeks gestation) and at very low birth weight(Hodel, 2018). Alterations include decreased prefrontal gray matter volume and surface area throughout childhood and adolescence. Decreased frontal white matter volume and poorer white matter integrity in frontal-lobe tracts have also been reported, suggesting that cortical development may be delayed or permanently impaired(Volpe, 2009). Specific deficits in executive functioning in preterm children, particularly in males have been documented across childhood and adolescence (Hodel, 2018; Nosarti et al., 2006).

Mid-to-late gestational brain development includes neurogenesis, migration, dendrite arborization and synaptogenesis, which are heavily dependent on GABA and glutamatergic neural systems and are at risk of disruption in the extra-uterine environment after preterm birth (Ben-Ari, 2006, 2014; Represa & Ben-Ari, 2005). Both neurons and astrocyte participate to the metabolism of GABA and glutamate as well as their transport and signaling (Ishibashi, Egawa, & Fukuda, 2019). Consistent with observation, ex-vivo studies of the preterm brain have demonstrated specific loss of GABAergic neuron populations in males (Lacaille et al., 2019), alteration of GABA receptor subunits and disorganized neuronal migration and differentiation (Robinson, Li, Dechant, & Cohen, 2006; Shaw, Palliser, Dyson, Berry, & Hirst, 2018; Shaw, Palliser, Walker, & Hirst, 2015).

Recent magnetic resonance spectroscopy imaging studies have suggested that GABA concentrations may be lower in preterm than term infants, but cannot assess the components of the GABAergic system that may have been altered during development. Recent reports have shown lower GABA and glutamate concentrations at term-equivalent age and a negative correlation with functional connectivity in small cohorts of preterm infants (born <35 weeks gestational age) compared to healthy full-term infants (Kwon et al., 2014; Tanifuji et al., 2017; Tomiyasu et al., 2017). Additionally, GABA concentrations were shown to correlate negatively with increasing gestational age at birth and were lower in the preterm infant brain at term equivalent age compared with healthy term controls and older children (6–16 years) (Basu et al., 2020). Similar imbalance between neuronal excitation and inhibition in the prefrontal cortex (PFC) (Marín, 2012) and decrease in prefrontal GABA concentrations have been associated with a number of psychiatric disorders, including autism spectrum disorder (ASD)(Schür et al., 2016), known to affect a high proportion of prematurely born surviving adults(Adams-Chapman & Stoll, 2006; Arpino et al., 2010; Knuesel et al., 2014; Marín, 2012; Meldrum et al., 2013; Ream & Lehwald, 2018).

Despite emerging links between psychiatric disorders and prematurity, the specific impact of preterm birth on the early maturation of neurotransmitter systems needs to be determined and compared between sexes. Thorough assessment of maturation assessment of the GABAergic system, which comprises the proteins necessary for GABA biosynthesis, degradation, release, signaling and reuptake by neurons and astrocytes, in the developing human frontal cortex is necessary to understand how prematurity results in psychiatric disorders in the absence of visible lesion and why male infants are more susceptible to develop such pathologies.

Here we delineate the molecular development of the GABAergic system in the dorsolateral PFC (Brodmann area 10; BA10), which plays a role in attention and working memory (Wang et al., 2018) and is altered in preterm infants and cases of psychiatric disorders(Hodel, 2018). Using an array of genes, we first assessed the maturation state of the GABAergic system from 0 to 8 months in human preterm post-mortem brain samples. We then showed that the development of the GABAergic system is delayed in male preterm infants, with major alterations on the astrocytic function. Taken together, this study shows that the maturation of GABAergic system is more dynamic in male infants potentially rendering them more susceptible than females to perinatal insults.

## Materials and Methods

### Human Samples

Human samples were obtained from the NIH Neurobiobank at University of Maryland, Baltimore, MD (ID #709). BA10 sections from 17 male and 9 female term infants ranging from 0 to 8 month were obtained to assess the normal maturation of the system at perinatal ages (i.e. cross sectional study, Table 1A,B). Additionally, sections from 7 male and 6 female preterm infants were obtained to compare their development to age-matched term infants (i.e. comparative study, Table 1 A,C). Mean age of death (absolute age) was 1.5 months in term infants and 3.7 months for preterm infants delivered between 26 and 34 weeks of gestation (average corrected age 1.1 months). Sex, completed gestational weeks and cause of death varied, but the majority were due to sudden infant death syndrome and none were attributed to CNS infection, hemorrhage or malformation (Table 1B,C). Genetic diseases or anatomic birth defects were excluded. Brodmann Area10 (BA10) formalin-fixed brain samples were cut into 0.5-cm-thick coronal slices and preserved in 10% neutral buffered formalin; matched frozen tissues were preserved at −80C.

**Table 1.**
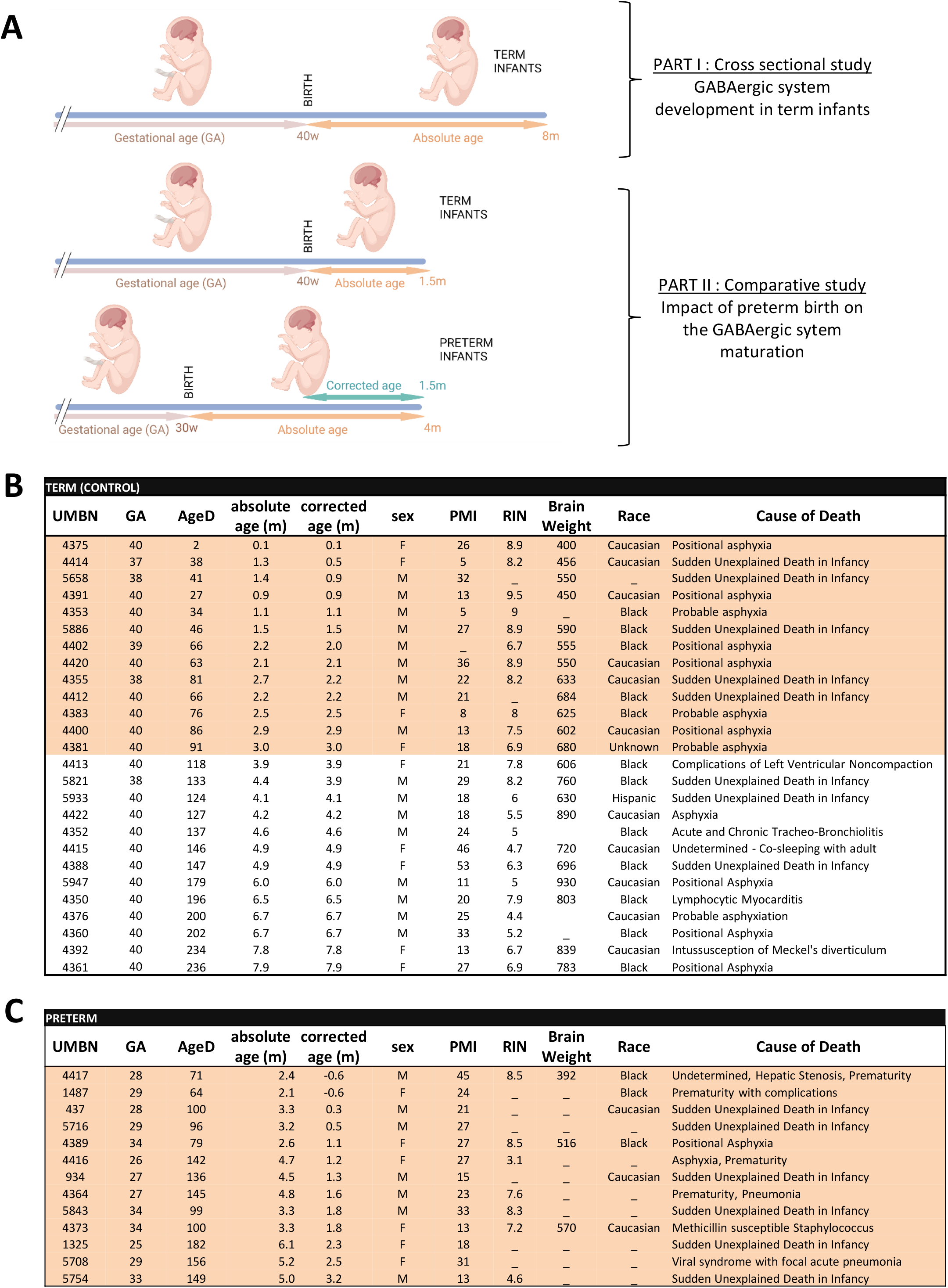
Experimental design and donor information. **A** Schematic showing the experimental design and the age terminology (Created with BioRender.com). List of donors (from NIH NeuroBioBank, University of Maryland, Baltimore, MD). **B** Term and **C** preterm, male and female infants. All the term donors were included in the cross sectional study. Donors highlighted in light orange were included in the comparative study. AgeD, Age at death in days; GA, gestational age; m, months; PMI, postmortem interval; RIN, RNA integrity number; UMBN, University of Maryland Bank Number.

### Real-Time PCR (RT-PCR)

BA10 tissues were homogenized in TRIzol™ Reagent (ThermoFisher, Waltham, MA, USA, 15596018); total RNA was extracted with the RNeasy Mini Kit (Qiagen Venlo, Netherlands, 74104) and quantified with a Nanodrop ND-2000C (ThermoFisher). 1 μg of RNA was used to make cDNA with the iScript cDNA Synthesis Kit (Bio-Rad, 1708891). All primer pairs were designed and validated in-house for efficiency and specificity. RT-PCR experiments were performed on cDNA samples in presence of SsoAdvanced Universal SYBR Green Supermix (Bio-Rad, Hercules, CA, USA, 1725271) with specific primers at 100 nM using the ABI Prism 7500 Sequence Detection System (ThermoFisher). The cDNA-generated signals for target genes were normalized with transferrin receptor protein 1 (*tfrc*). The regulation was determined with the 2^-ΔΔCq^ method. Results are expressed as fold change (FC) to the control group.

### Western Blot

Human samples were homogenized in RIPA lysis buffer with proteinase inhibitors (Santa Cruz Biotechnology, Dallas, TX, USA, sc24948). Protein extracts, 40 μg per lane, were loaded onto 4–20% gradient gels (NuSep Inc, Germantown, MD, USA, NB10-420). Gels were electrotransferred to a 0.2 μm nitrocellulose membrane (Bio-Rad, 1620174). Blots were blocked in 5% milk in TBST for 1 h, and then incubated at 4 °C overnight with one of the following antibodies raised against: AATM (1:1000, sc-271702, Santa Cruz Biotechnology), AQP4 (1:1000, sc-390488), CX30 (1:1000, sc-81802), CX43 (1:1000, sc-271837), EAAT1 (1:500, sc-515839), GABRA1-6 (1:250, sc-376282), GAPDH (1:2000, sc-32233, GAT-3 (1:250, sc-376001), GDH1/2 (1:1000, sc-515542), GEPH (1:1000, sc-25311), GS (1:1000, sc-74430), MAOB (1:250, sc-515354), SNAT5 (1:1000, sc-515813), VIAAT (1:500, sc-393373), GBRD (1:250, Novus Biologicals, Littleton, CO, USA. NB300-200) or GABRG1 (1:100, Alomone labs, Jerusalem, Israel, AGA-016). Bands were detected with appropriate horseradish peroxide-conjugated secondary antibodies, reacted with chemiluminescent ECL substrate (Bio-Rad, 1705060) and visualized with a BioRad ChemiDoc Imaging system. Band intensity was measured using the ImageJ program (NIH) and normalized with GAPDH.

### Immunohistochemical procedure

Formalin-fixed tissues were cryoprotected in a 30% sucrose solution and embedded in Tissue-Tek® O.C.T. Compound (Sakura® Finetek, Torrance, CA, USA). Blocks were cut into 25-μm-thick sections on a cryostat and mounted on Superfrost Plus (ThermoFisher) glass slides. Frozen sections were allowed to equilibrate to room temperature for 2 hours before staining.

#### Procedure

Tissue sections were rinsed in PBS-Triton 0.3% (PBS-T) then blocked in PBS-T with 10% normal donkey serum (NDS) followed by overnight incubation at 4°C in PBS-T-10% NDS with primary antibodies raised against: Calbindin (1:1000, Swant, Marly, Switzerland, CB300 or CB38), Calretinin (1:1000, Millipore Sigma, AB1550), Gad65-67 (1:200 Santa Cruz Biotechnology, sc-365180), GFAP (1:1000, Agilent Dako, Glostrup, Denmark, Z0334), Neuropeptide Y (1:500, Immunostar, Hudson, WI, USA, 22940) or Somatostatin (1:300, Santa Cruz Biotechnology, sc7819). For secondary detection, appropriately matched Alexa Fluor-conjugated secondary antibodies (1:500, ThermoFisher) were incubated 90 min in PBS-T at room temperature. Sections were incubated with DAPI, mounted in Fluoromount G (ThermoFisher) and coverslipped before epifluorescence examination with a slide scanner (Leica DMI6000 B, Leica, Wetzlar, Germany).

#### Quantification

Cell density was assessed in the upper layers (UL), lower layers (LL) and the subcortical white matter (SC-WM) of BA10 and expressed in cells per mm^2^. Cortical layering was determined with DAPI counterstaining. Cell quantification was performed using the Qpath software (Edinburgh, UK). All counts were performed blind to condition.

### Statistics

All experiments and analyses were performed blind to conditions. Statistical analysis was performed using PRISM software (GraphPad Software 6.0, San Diego, CA, USA). Normal distribution of each dataset was analyzed by Shapiro-Wilk test. When two conditions were compared, data were analyzed with a nonparametric Mann-Whitney test. When four experimental groups were assessed and two variables were taken into consideration, data were analyzed with a two-way ANOVA with Fisher LSD, Sidak’s or Tukey’s multiple comparisons. The maturation index and the developmental trajectories of transcripts and proteins in term infants was quantified by fitting linear regression slopes to measure development through postnatal months. To compare the difference of two slopes, data were analyzed with an ANCOVA The null hypothesis was rejected for alpha greater than 5%.

## Results

BA10, also known as dorsolateral prefrontal cortex, is one of the brain regions most frequently altered in psychiatric diseases. To address whether prematurity alters the maturation of the GABAergic system, a cross sectional developmental study was designed. The expression of 47 transcripts related to the GABA system were determined in pathology specimens from male and female term infant across multiple early postnatal time points (Table 1A,B; Fig. 1A-D). Screened genes encoding different components of the GABAergic signaling system included: GABA receptor subunits, metabolic enzymes, transporters, ion channels associated to the GABAergic function, interneuron specific markers, GABA receptor anchorage proteins and enzymes involved in the metabolism of glutamate and GABA (Table 2).

**Figure 1.**
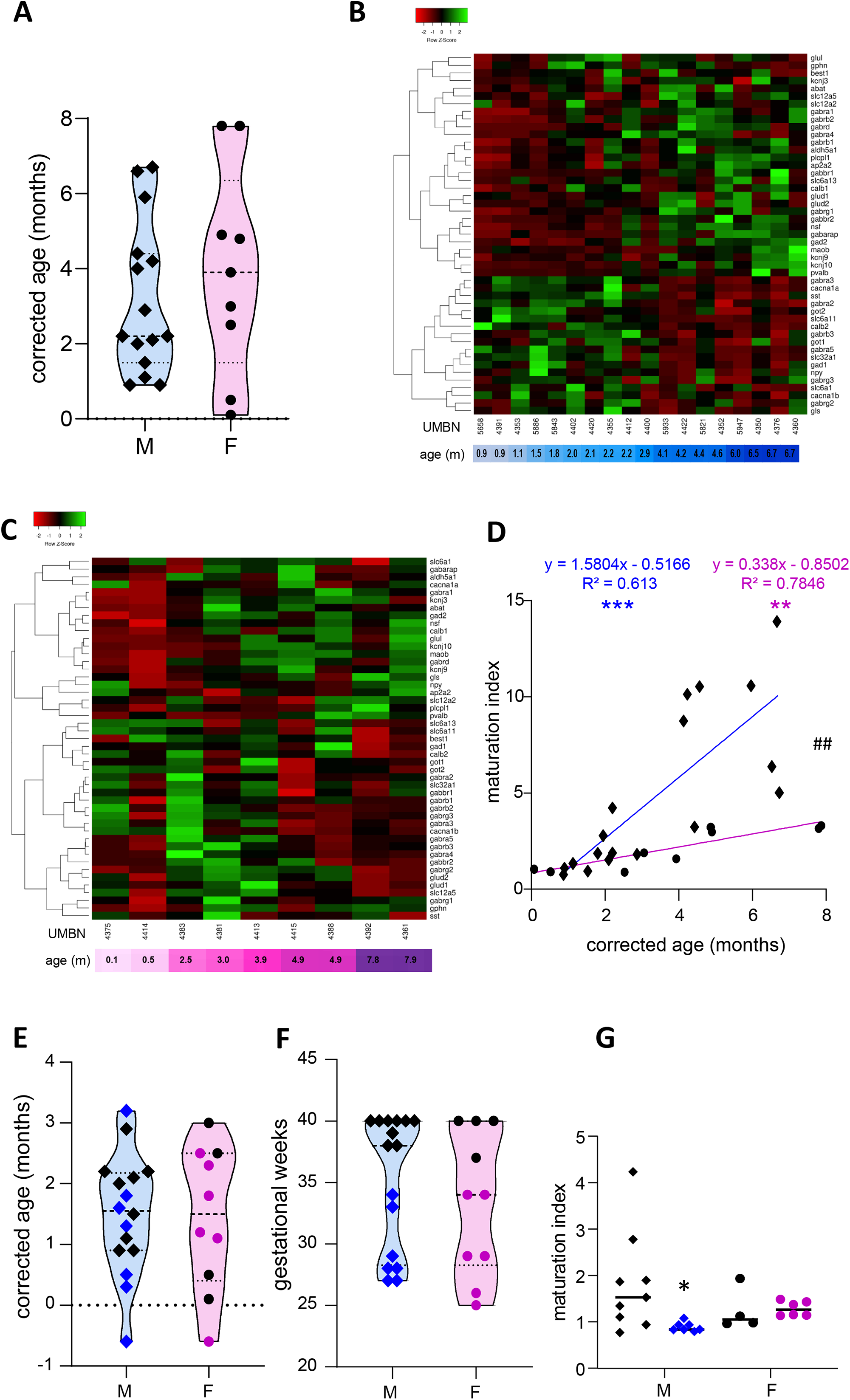
Preterm birth alters the maturation of the GABAergic system. **A** Violin plot displaying the distribution of male (M, blue, diamonds) and female (F, pink, circles) term infants included in the cross sectional study. Violin plots show individual values and include minima, maxima and median values. **B-C**. Heatmap of GABA-associated gene expression means in the BA10 of **B** male and **C** female term infants, arranged using hierarchical clustering (Complete linkage with Pearson distance measurement method). **D**. Maturation index based on the heatmap clustering of male (blue, diamonds) and female (purple, circles) term infants cross sectional study, modeled with a linear regression, **p < 0.01, **p <0.001. ##p <0.01 (ANCOVA, Analysis of Co-variance). **E-F** Violin plot displaying the distribution of male (M, blue, diamonds) and female (F, purple, circles) infants, term (black) versus preterm (colored) infants included in the comparative study according to (**E**) their corrected age (in months) and (**F**) gestational weeks. **F** Maturation index of male and female term (black) and preterm (colored) infants comparative study, based on the heatmap clustering of the term infants cross-sectional study. *p<0.05 (two-way ANOVA with Sidak’s multiple comparisons).

**Table 2.**
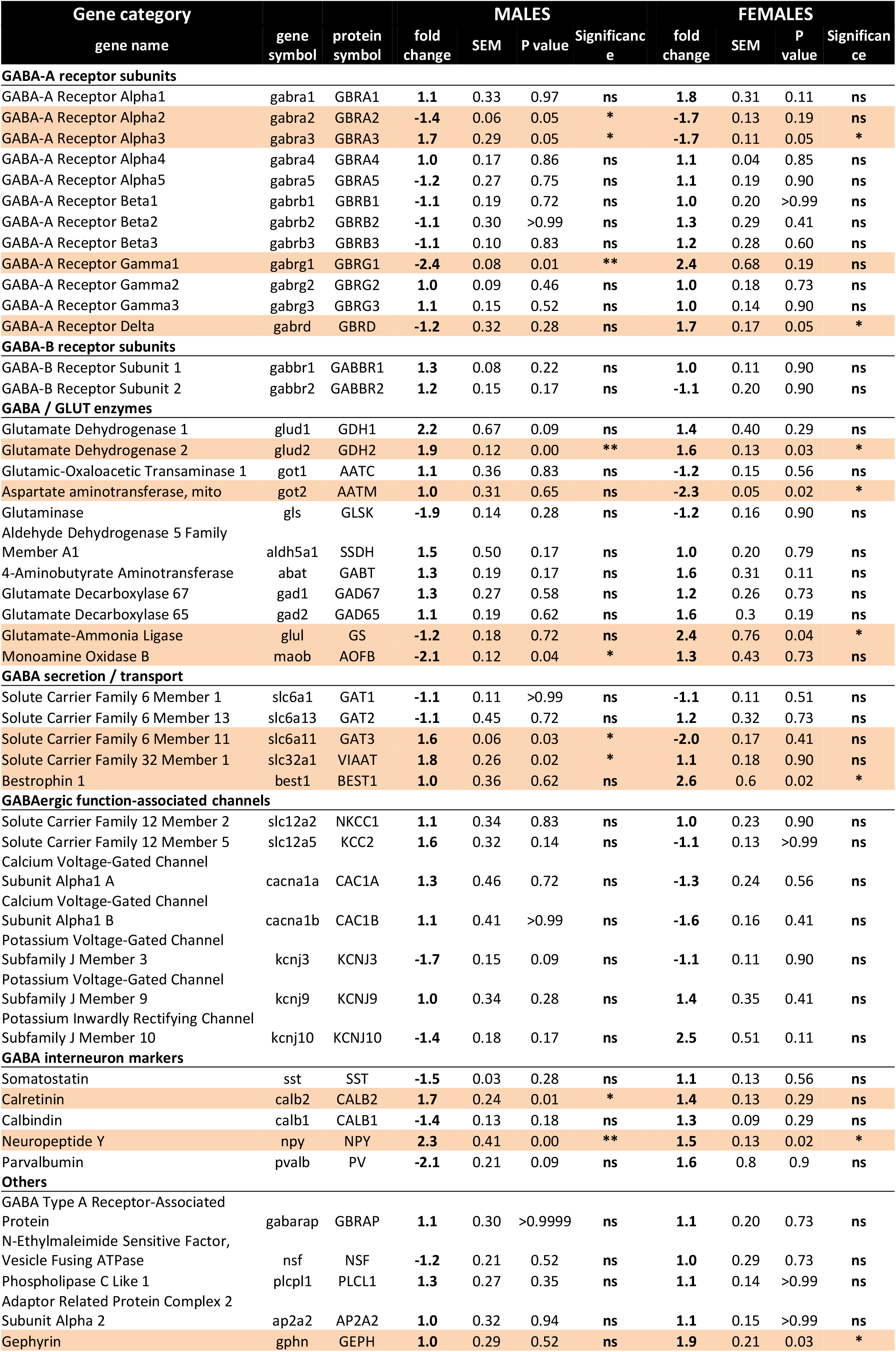

### The maturation of the GABAergic system is more dynamic in male than female term infants

To estimate the overall maturation of the GABAergic system based on gene expression level, a maturation index was calculated as previously described(Gandal, Nesbitt, McCurdy, & Alter, 2012). Expression levels for most genes changed monotonically across development, either increasing or decreasing. The ratio of the average of developmentally upregulated over downregulated genes also increase monotonically with time. The genes analyzed were plotted into a heatmap and separated into up and down-regulated genes using the first linkage of the heatmap for male (Fig. 1B) and female infants independently (Fig. 1C). Equal weight was given to all genes within the maturation index. The ratio of averaged upregulated (29 genes) over averaged downregulated genes (18 genes) was used as a maturation index for the development of the GABAergic system in term males; and similarly in term female infants with 24 upregulated and 23 downregulated genes (Fig. 1D). As expected, the maturation index significantly correlated with the age of the donor in males (R^2^ = 0.61 p<0.0001, Fig. 1D) and in females (R^2^ = 0.78 p<0.0001, Fig. 1D). Notably, the slope of the maturation index was significantly greater in males compared to females (p<0.001, Fig. 1D), indicating broader maturation dynamic in the former.

### The maturation of the GABAergic system is altered in preterm infants

To investigate the possibility of a maturation delay of the GABAergic system in preterm infants, a comparative study was designed. Corrected age matched preterm infants and the youngest term infants (0 to 3 months) were compared (Table 1A,C Fig.; Fig. 1E-G). The same transcripts were analyzed in male and female term and preterm cerebral cortices. The maturation index of term and preterm infants was calculated as ratio of the average of developmentally upregulated over downregulated genes, as previously defined with the cross-sectional developmental study. Interestingly, the maturation index was decreased in male (–50% p<0.05, Fig. 1G) but not female preterm infants (–6% ns, Fig. 1G) when compared respectively to their matched term group. The reduced maturation index in preterm male infants signifies that prematurity induces gene expression changes in the opposite direction from that occurring normally during GABAergic system development. To gain insight into the function of these genes, their expression was analyzed individually (Table 2).

### Preterm birth delays the development of the GABAergic system in male infants

Out of 47 transcripts, nine were significantly dysregulated in male preterm infants. Only *gabra2* varied according to its predicted developmental trajectory by linear regression, which suggest an accelerated maturation. Indeed, *gabra2* expression is predicted to decrease over time (p<0.01, Fig. 2A) and was decreased in preterm male infants (FC= −1.4 p<0.5, Table 2; Fig. 2A’). Most of the transcripts varied oppositely to their predicted developmental trajectory suggesting an overall developmental delay (Table 2; Fig. 2). The dysregulated transcripts included: two GABAA receptor subunits, *gabra3* (FC= 1.7 p<0.05, Table 2) and *gabrag1* (FC= −2.4 p<0.01, Table S2; Fig. 2B,B’); two enzymes: *glud2* (enzyme catalyzing the reversible interconversion of glutamate to α-ketoglutarate, FC= 1.9 p<0.01, Table 2; Fig. 2C-C’) and *maob* (enzyme responsible for the production of GABA in astrocytes, FC= −2.1 p<0.05, Table 2; Fig. 2D,D’); two transporters: *slc6a11* (gaba transporter in astrocytes, FC= 1.6 p<0.05, Table 2; Fig. 2E,E’) and *slc32a1* (vesicular transporter, FC= 1.6 p<0.05, Table 2); two markers of GABAergic interneuron subpopulation: *crt* (calretinin, FC= 1.7 p<0.05, Table 2) and *npy* (neuropeptide Y, FC= 2.3 p<0.001, Table 2). Further examination revealed that four of these genes were associated to the astrocytic functions including *gabrg1, gdh2, maob* and *gat3*. Additional astrocyte-related genes were trending toward significance (Table 2; *glud1* FC= 2.2, *gls* FC= −1.9, and *aldh5a1* FC= 1.5, enzymes part of the TCA cycle). The majority of protein expression levels assessed by western blot covaried positively with the mRNA changes in males born preterm, suggesting that the mRNAs of interest were actively synthetized (Fig. 2).

**Figure 2.**
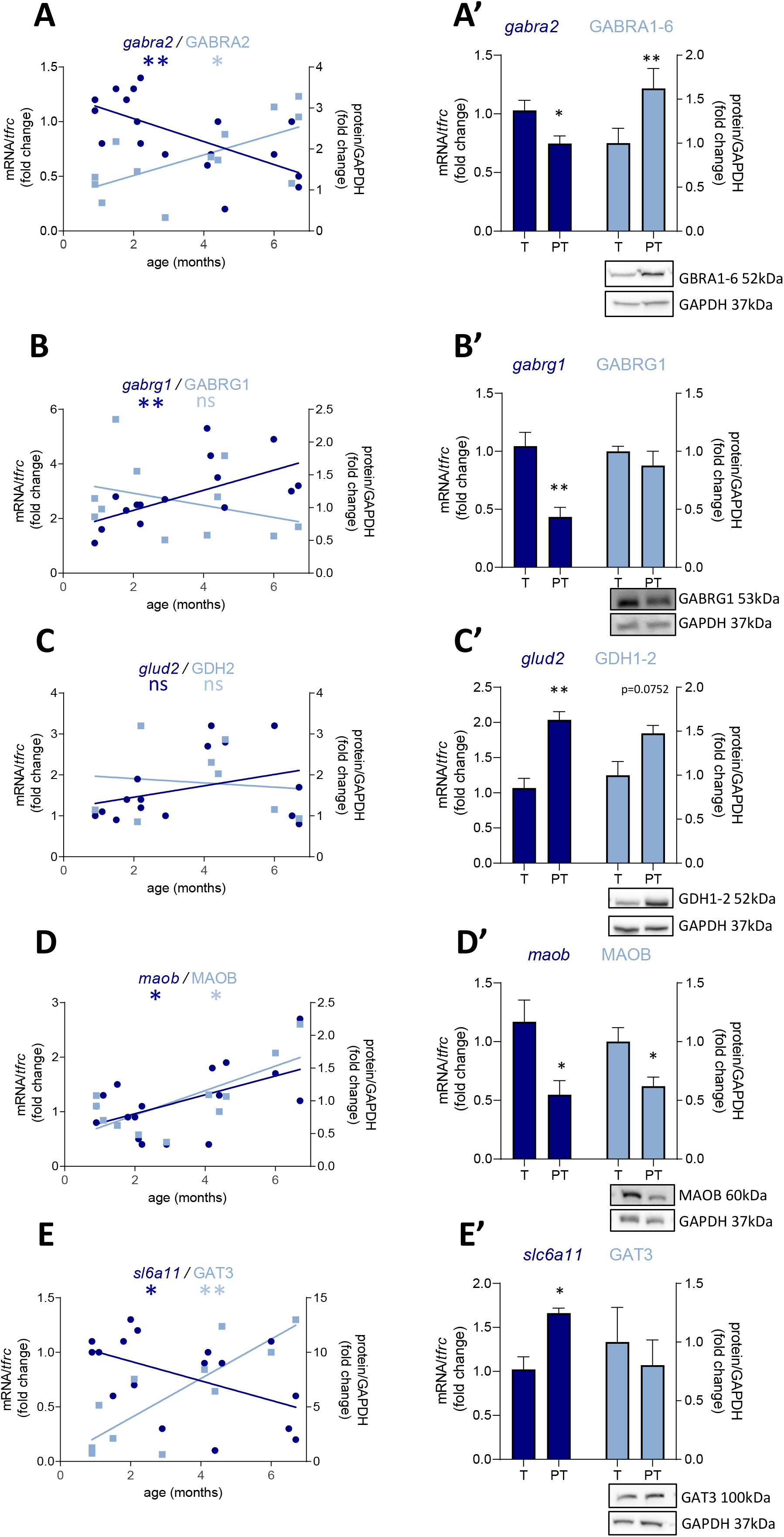
Preterm birth alters the expression of GABA-related transcripts (dark blue) and proteins (light blue) in male infant BA10. **A-E** Developmental regulation in term infants included in the cross sectional study, modeled with a linear regression *p < 0.05, **p <0.01; left column. **A’-E’** Effect of preterm birth (comparative study) on the expression of GABA-related transcripts and proteins. Quantification of mRNA level changes by qRT-PCR of **A-A’** *gabra2*, **B-B’** *gabrg1*, **C-C’** *glud2*, **D-D** *maob* and **E-E’** *slc6a11. tfrc* was used for normalization. Quantification of protein expression changes by Western blot of **A-A’** GABRA1-6, **B-B’** GABRG1, **C-C’** GDH2, **D-D’** MAOB and **E-E’** GAT3. GAPDH was used for normalization. *p < 0.05, **p <0.01 (Mann–Whitney). Representative blot below the graph. PT, preterm; T, term.

### Preterm birth accelerates the development of the GABAergic system in female infants

In female preterm infants, 8 out of 47 transcripts were significantly dysregulated, most varied accordingly to their developmental trajectory suggesting an accelerated development or to resemble their non-corrected age counterpart (Table 2; Fig. 3) -*gabra3* and *gabrd* (GABA_A_ receptor subunits) -*glud2, got2* and *glul* (enzymes responsible for the glutamine to glutamate conversion) –*best1* (GABA transporter in astrocytes) -*npy* (interneuron subtype) and *geph* (responsible for GABA receptor anchorage). Similarly to males, the majority of protein expression mostly covaried with mRNA in female preterm infants (Fig. 3).

**Figure 3.**
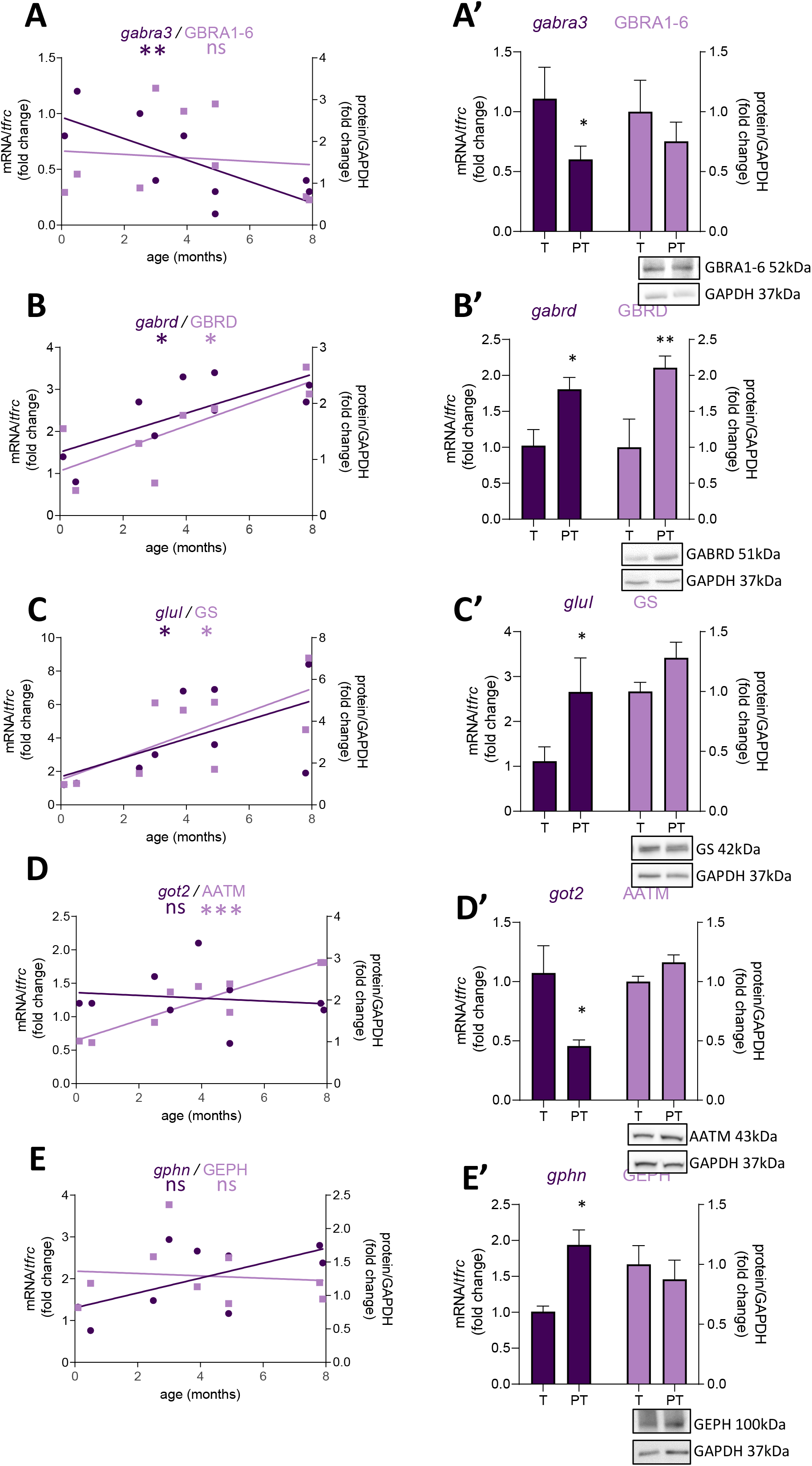
Preterm birth impacts the expression of GABA-related transcripts (dark purple) and proteins (light purple) in female infants. **A-E** Developmental regulation in term infants included in the cross sectional study, modeled with a non-linear regression *p < 0.05, **p <0.01, ***p <0.001; left column. **A’-E’** Effect of preterm birth on the expression of GABA-related transcripts and proteins (comparative study). Quantification of mRNA level changes by qRT-PCR of **A-A’** *gabra3*, **B-B’** *gabrd*, **C-C’** *glul*, **D-D’** *got2* and **E-E’** *gphn. tfrc* was used for normalization. Quantification of protein expression changes by Western blot of **A-A’** GABRA1-6, **B-B’** GABRD, **C-C’** GS, **D-D’** AATM and **E-E’** GEPHN. GAPDH was used for normalization. *p < 0.05, **p <0.01 (Mann–Whitney). Representative blot below the graph. PT, preterm; T, term.

### Preterm birth primarily affects the astrocytic GABA function in male infants

Since most of the perturbations observed in male preterm infants seemed to be related to the astrocytic regulation of GABAergic signaling, ten supplementary transcripts were analyzed to evaluate specifically the metabolism and morphogenesis and ten others to evaluate the involvement of astrocytes in the GABAergic system (Fig. 4; Table 3). A maturation index specific to astrocytes was calculated with the averaged upregulated over averaged downregulated genes using the first linkage of the astrocyte-specific heatmap, individually in males (Fig. 4A) and females (Fig. 4B). The genes plotted were 14 genes related to astrocytes and GABA included in Table 2 (*glul, best1, abat, aldh5a1, glud2, glud1, gabrg1, maob, kcnj10, slc6a11, got1, gabrg3, gls, got2*) and 20 newly added genes (Table 3). The maturation index was significantly correlated with the age of the donors both in males (R^2^ = 0.54 p<0.001, Fig 4C) and in females (R^2^ = 0.44 p<0.001, Fig 4D). Across the spam of ages examined (0 to 8 months), astrocytic maturation was significantly more dynamic in male infants since the slope of their maturation index was significantly superior to the one in female infants (p<0.01, Fig. 4C). The same transcripts were analyzed in term and preterm infants to estimate their astrocyte-related maturation indices. The astrocyte-maturation index was significantly decreased in male (–50% p<0.05, Fig. 4D) but not female preterm infants (−13% ns, Fig. 4D).

**Figure 4.**
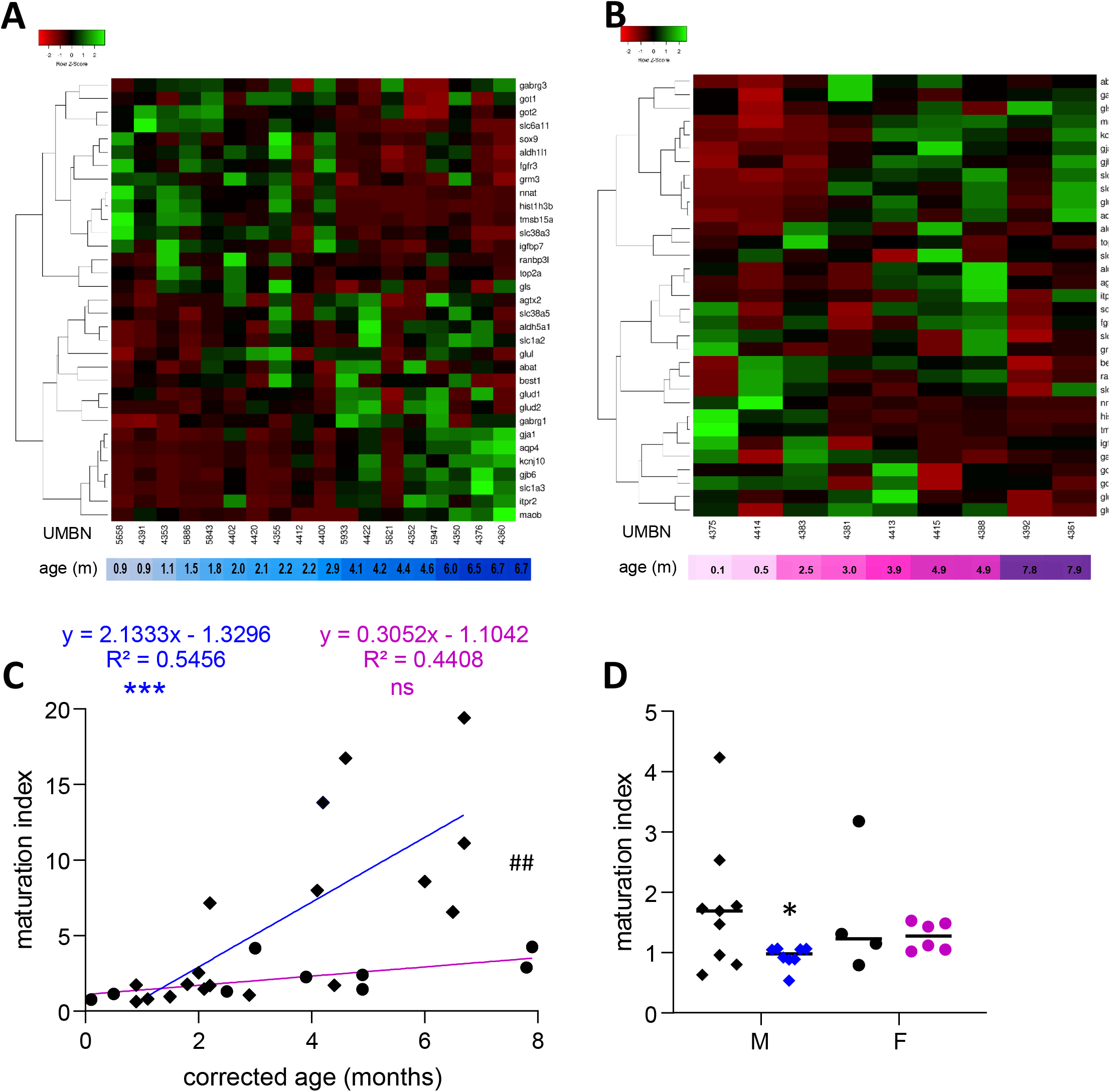
Preterm birth alters the maturation of the GABAergic system in astrocytes. **A** Heatmap of astrocyte-related gene expression means in the BA10 of **A** male and **B** female term included in the cross sectional study, arranged using hierarchical clustering (Complete linkage with Pearson distance measurement method). **C** Maturation index based on the heatmap clustering of male (blue, diamonds) and female (purple, circles) term infants cross sectional study, modeled with a linear regression ***p <0.001). ##p <0.01 (ANCOVA, Analysis of Co-variance). **D** Maturation index of male and female term (black) and preterm (colored) infants comparative study, based on the heatmap clustering of the term infants cross-sectional study. *p<0.05 (two-way ANOVA with Sidak’s multiple comparisons).

**Table 3.**
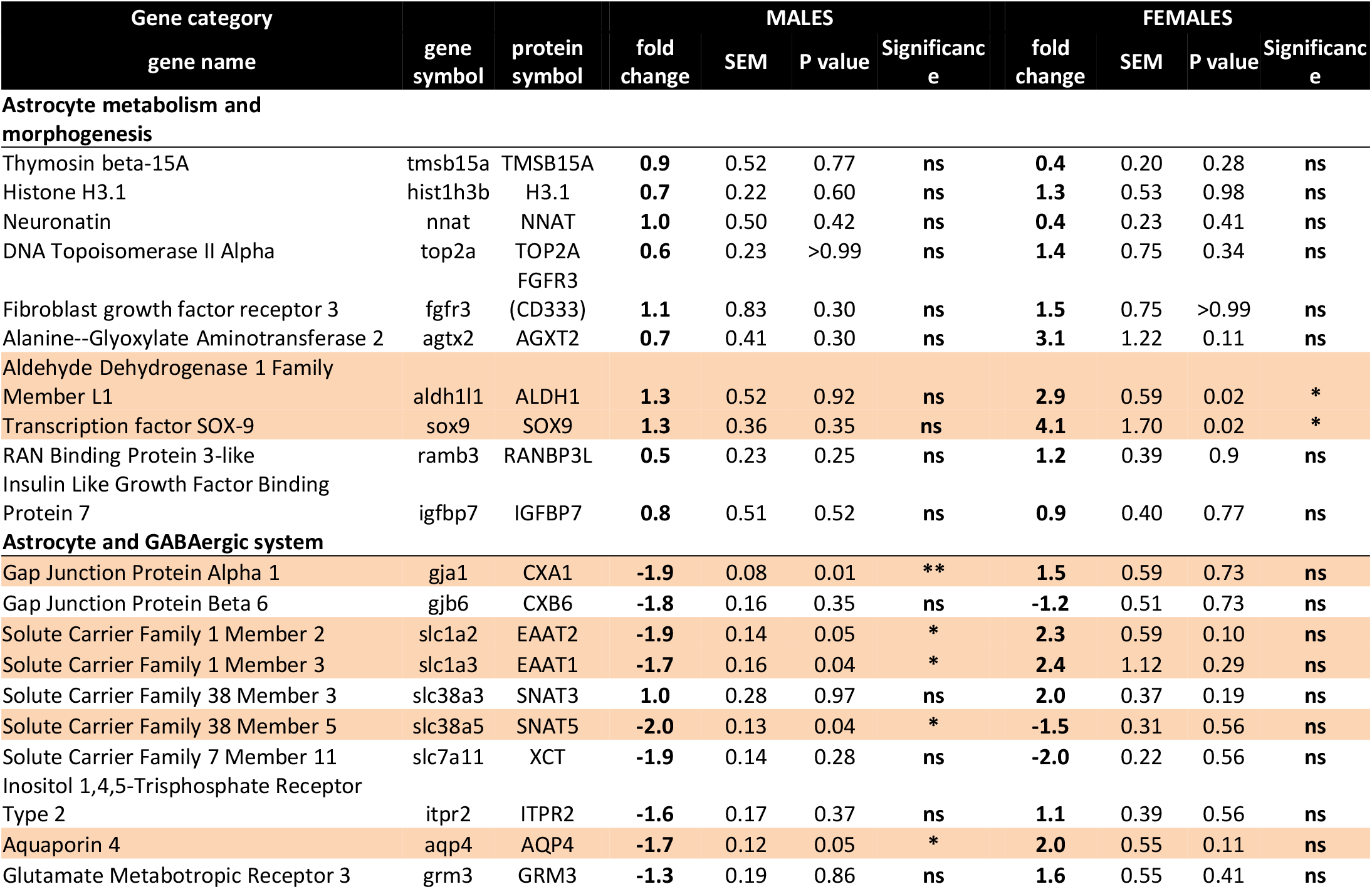

Out of the ten genes related to astrocyte metabolism and morphogenesis, two were up-regulated in females only, both linked to astrocyte intrinsic metabolism. -*aldh1l1*, (metabolic enzyme, FC= 2.9 p<0.05, Table 3) and –*sox9* (transcription factor, FC= 4.1 p<0.05, Table 3). Regarding the genes associated with the astrocytic GABA function, five were significantly regulated in males. In accordance with the decreased maturation index, these transcripts varied in the opposite way to their developmental trajectories, predicted by linear regression (Fig. 5). The dysregulated transcripts included: -*gja1* (gap junction allowing intercellular communication between astrocytes, FC= −1.9 p<0.01, Table 3; Fig. 5A) -*slc1a2* and *slc1a3* (glutamate transporters, FC= −1.9 p<0.05 and FC= −1.7 p<0.05, Table 3; Fig. 5B,C) -*slc38a5* (glutamine transporter, FC= −2.0 p<0.05, Table 3; Fig. 5D) and *aqp4* (water channel, FC= −1.7 p<0.05; Table 3; Fig. 5E). Protein and mRNA expression levels followed the same pattern in preterm infants samples (Fig. 5).

**Figure 5.**
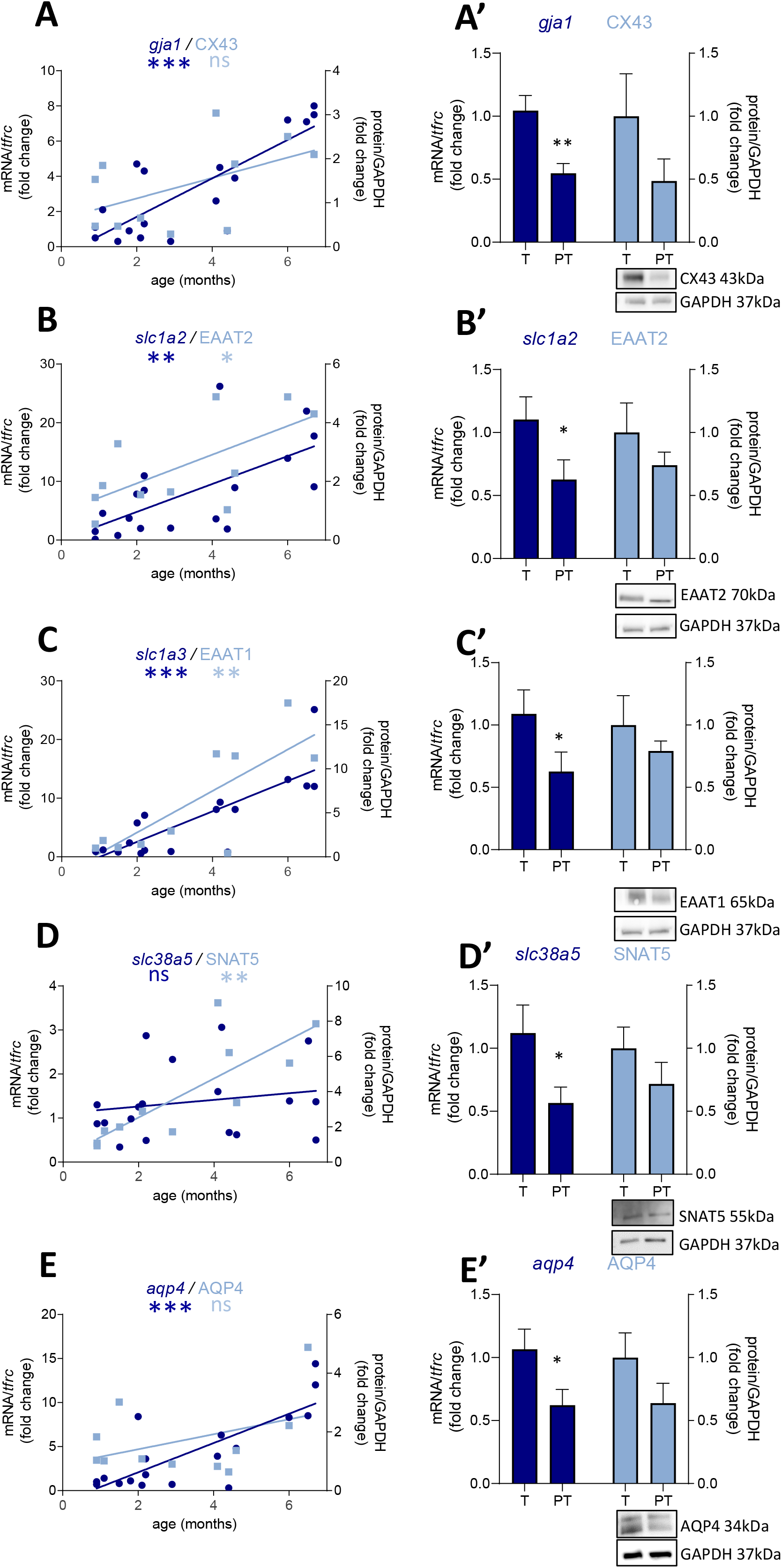
Preterm birth impacts the expression of astrocyte-related transcripts (dark blue) and proteins (light blue) in the BA10 of males infants. **A-E** Developmental regulation in term infants included in the cross sectional study, modeled with a non-linear regression (*p < 0.05, **p <0.01, ***p <0.001; left column). **A’-E’** Effect of preterm birth on GABA-related transcripts and proteins (comparative study). Quantification of mRNA level changes by qRT-PCR of **A-A’** *gja1*, **B-B’** *slc1a2*, **C-C’** *slc1a3*, **D-D**, *slc38a5* and **E-E’** *aqp4. tfrc* was used for normalization. Quantification of protein expression changes by Western blot of **A-A’** CX43, **B-B’** EAAT2, **C-C’** EAAT1, **D-D’** SNAT5 and **E-E’** AQP4. GAPDH was used for normalization. *p < 0.05, **p <0.01 (Mann–Whitney). Representative blot below the graph. PT, preterm; T, term.

### The effect of preterm birth on the maturation of the GABAergic system is not due to a change in the number of astrocytes

To address whether the observed molecular changes were due to a reduced number of cells or to a cellular maturation defect, both GABA-producing cell (interneurons) and astrocyte densities were assessed in three subdivisions of BA10: the upper layers (ULs), lower layers (LLs) and the sub-cortical white matter (SC-WM). The density of major subclasses of cortical interneurons was assessed by immunostaining for GAD65-67, SST, CLB, CRT and NPY (Fig. 6). GAD65-67-positive cell density was significantly decreased in the ULs of preterm male infants as compared with term male infants (UL –37% p<0.05, LL –28% ns, SC-WM -27% ns, Fig. 6A,A’). A significant reduction in the density of SST was observed in both UL and LL (UL –43% p<0.05, LL – 30% p<0.05, SC-WM -23% ns, Fig. 6B,B’) and of CLB interneurons in the UP but LL (UL –47% p<0.05, LL -17% ns, SCWM: -1% ns, Fig. 6C,C’). No change was observed in the density of CRT or NPY interneurons (Fig. 6D,E). PV cells were not detected in this set of human samples, although these cells could be detected at later developmental stages. No statistically significant change in total cortical layer widths was discernable based on Nissl staining (data not shown), suggesting overall preservation of pyramidal cells at this age. No difference was observed in female infants, but statistical power was limited by the small sample size (Fig. 6A’-E’).

**Figure 6.**
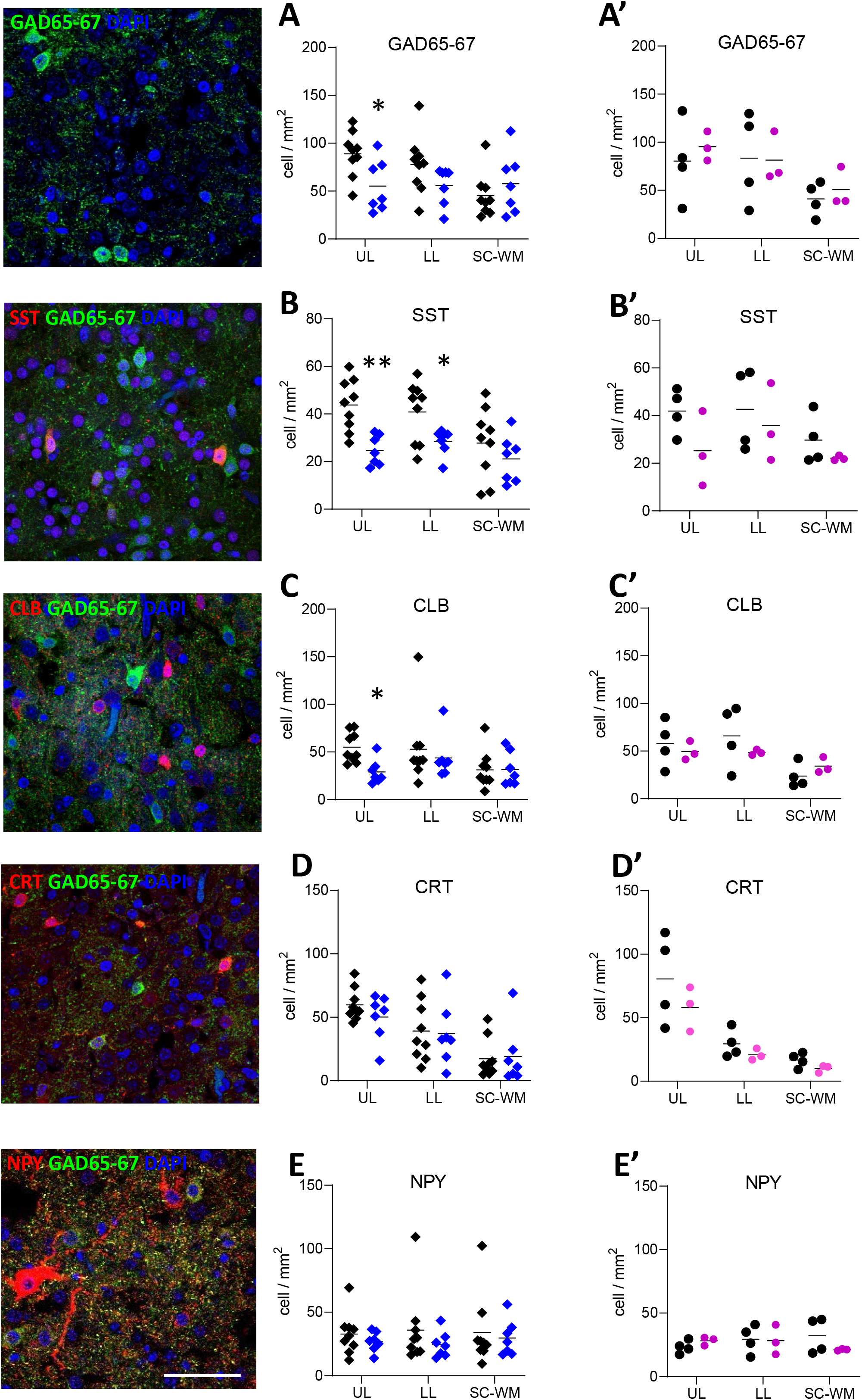
Preterm birth impacts interneuron density in the BA10. Illustrations of **A** glutamate decarboxylase 65 and 67 (GAD65-67, green), **B** SST (red), **C** CLB (red), **D** CRT (red), and **E** NPY (red) immunofluorescent staining. Scale bar 50 μm. Respective quantification of immmunopositive cells in the upper layers (UL), lower layers (LL) and subcortical white matter (SC-WM) in the BA10 of **A-E** male (blue, diamonds) and **A’-E’** female (pink, circles) term (black) versus preterm (colored) infants included in the comparative study. Scatter dot plots show the mean and individual dispersion of term (colored circles) and preterm infants (black circles); *p <0.05, **p<0.01 (two-way ANOVAs were performed followed by Fisher’s LSD tests for post hoc comparisons).

GFAP-positive cell density was not significantly affected by preterm birth in males (UL –14% ns, LL –62% ns, SC-WM -15% ns, Fig. 7A) and in females (UL –41% ns, LL – 25% ns, SC-WM -9% ns, Fig. 7A’). In accordance with the cell counts, GFAP mRNA and protein levels were not altered in premature infants (Fig. 7B-B’,C-C’). Notably, the expression of *gfap* represented by a linear regression, increased in the postnatal period in males, confirming the dynamic aspect of the system being broader than in females (p<0.05, Fig. 7B; ns, Fig. 7C).

**Figure 7.**
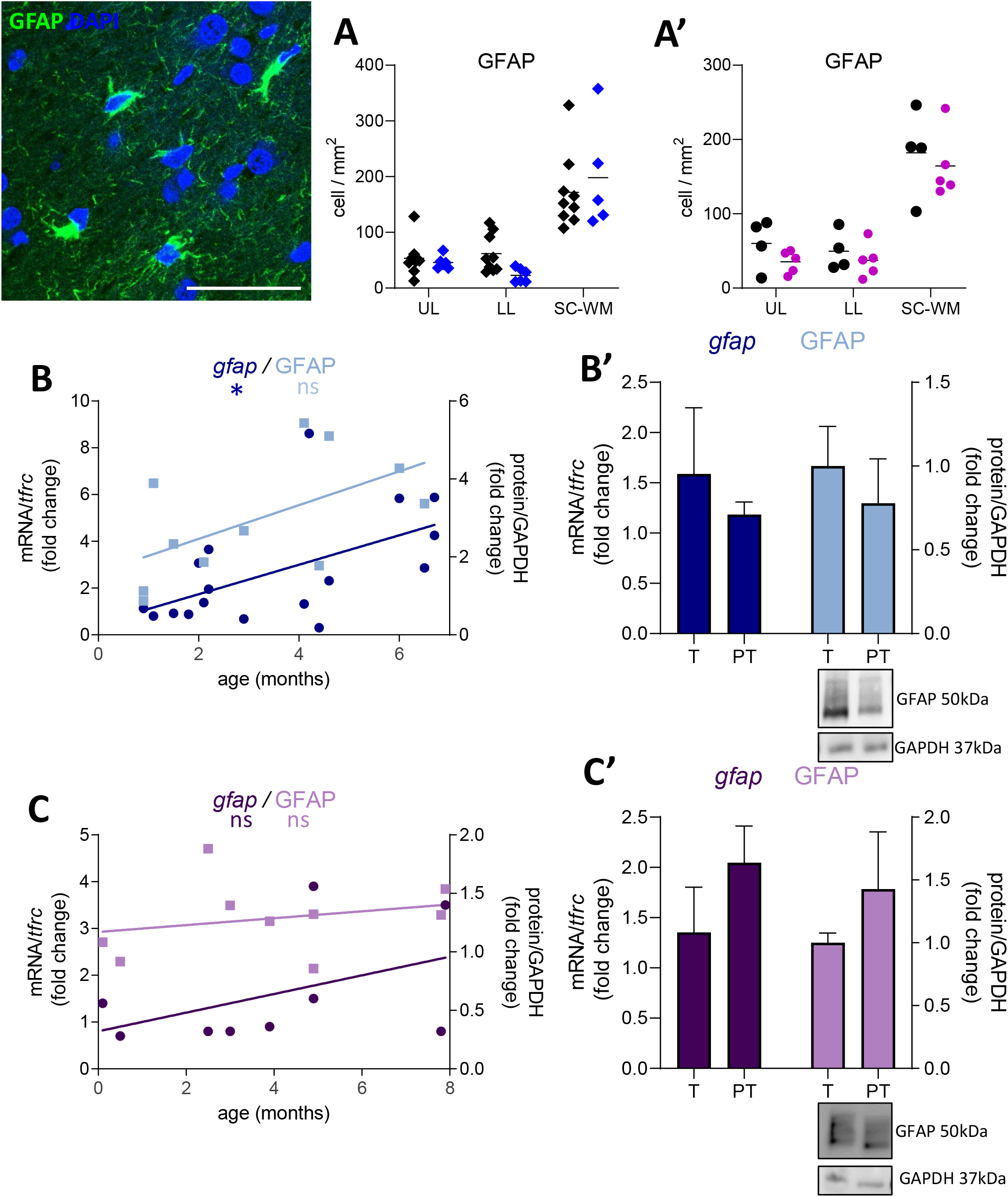
Effect of preterm birth on BA10 GFAP expression. Illustrations of GFAP-immunofluorescent staining. Scale bar X μm. **A-A’** Quantification of GFAP positive cells in the upper layers (UL), lower layers (LL) and subcortical white matter (SC-WM) in the BA10 of **A** male (blue, diamonds) and **A’** female (pink, circles) term (black) versus preterm (colored) infants included in the comparative study. Scatter dot plots show the mean and individual dispersion of term (black circles) and preterm infants (colored circles); (two-way ANOVAs were performed followed by Fisher’s LSD tests for post hoc comparisons). **B-C** Effect of preterm birth on the development of GFAP transcripts (dark) and proteins (light) in the BA10 of **B** male (blue) and **C** female (purple) infants, included in the cross sectional study, modeled with a non-linear regression *p < 0.05; left column. **B’-C’** Effect of preterm birth on GFAP expression (comparative study). Quantification of mRNA level changes by qRT-PCR of *gfap. tfrc* was used for normalization. Quantification of protein expression changes by Western blot of GFAP. GAPDH was used for normalization (Mann–Whitney). Representative blot below the graph. PT, preterm; T, term.

## Discussion

Understanding the effects of perinatal insults on GABAergic system development in male and female infants is critical to elucidating the mechanistic role it plays in the pathogenesis of neuropsychiatric disorders as well as the greater susceptibility of male infants to these pathologies. To assess the development of the GABAergic system, 47 genes related to GABA were analyzed in a cross-sectional study, in the BA10 (dorsolateral prefrontal cortex) of young male and female term infants (0-to 8-month-old). A maturation index was calculated based on the ratio of up-regulated versus down-regulated genes, as previously described (Gandal et al., 2012). The index was correlated with the age of the donor and was more dynamic in male than in female infants, as suggested by the steeper slope of the linear regression. Prior studies have shown that developmentally regulated differences in gene- and exon level expression exist between male and female brains in specific regions, including the PFC, and can have long lasting effects on brain development and plasticity (Mottron et al., 2015). Specifically, males are more susceptible than females to perturbations in genes involved in synaptic plasticity (Mottron et al., 2015). The flexibility of neuronal networks depends on both excitatory and inhibitory synapse plasticity, which notably relies on the number of postsynaptic GABAergic receptors (Barberis, 2020). Such GABAergic plasticity, elicited by the ventral hippocampus and basolateral amygdala, has been demonstrated in the prefrontal cortex (Caballero, Thomases, Flores-Barrera, Cass, & Tseng, 2014). Within a local network, the GABAergic system can also adapt its efficacy to the overall activity of the network, being upregulated in hyperactive networks and downregulated under conditions of reduced activity. This homeostatic plasticity is mediated by multiple pre- and postsynaptic mechanisms, including the concentration of GABA itself, regulated by synthetizing enzymes and transporters (Roth & Draguhn, 2012).

To evaluate the impact of premature birth on GABAergic system development, the prefrontal cortices from one-month-old term and one-month corrected-age very preterm infants were compared, using the same gene list. The calculation of the maturation index showed a decrease in male but not in female preterm infants, suggesting a maturation delay in the formers. The basal higher dynamicity in male infants could explain this difference, rendering the genes associated to the GABAergic system more susceptible to perturbation. The maturation of the GABAergic system could either be delayed to compensate for the immature excitatory system, or be the result of various noxious events suffered by preterm infants.

Out of the 47 genes analyzed, the differentially expressed genes (DEGs) affected by preterm birth were distinct between male and females. Male DEGs varied in opposition to their predicted developmental trajectory resulting in the overall developmental delay displayed by the maturation index. Closely looking at individual DEGs revealed that most of the gene regulation in male were tied to the astrocytic GABAergic function. For instance, prematurity markedly reduced the expression of two GABA_A_ receptor subunits: *gabra2*, enriched is astrocytes(Zhang et al., 2016), and *gabrg1*, which is specifically expressed by astrocytes(Batiuk et al., 2020). In these cells, GABA is depolarizing and contributes to maintaining GABAergic neuronal transmission via Cl^−^ efflux, buffering the Cl^−^ concentration of the GABAergic synapse(Isomura et al., 2003). Astrocytes also control the biosynthesis and turnover of GABA, both affected by prematurity. *Maob*, responsible for the synthesis of GABA from putrescine in astrocytes (Yoon et al., 2014), was downregulated while *gdh2*, an enzyme in charge of the turnover of glutamate, which catabolism is tied with GABA through the tricarboxylic acid cycle(Hertz, Wu, & Schousboe, 1978). Our findings suggest that this decreased synthesis and increased turnover would lead to a decrease of total GABA content in preterm male infants. A recently published study demonstrated that GABA concentration in the prefrontal cortex was lower in preterm infants when measured in vivo(Basu et al., 2020). Lower levels of brain GABA and glutamate concentrations have been associated with a variety of neurologic disorders including epilepsy, ASD and ADHD (Schür et al., 2016), which are more common in surviving premature infants (Ream & Lehwald, 2018).

In female preterm infants, the DEGs varied accordingly to their developmental trajectory suggesting either an accelerated development or compensatory mechanisms to resemble their non-corrected age counterpart. Preterm birth is a leading cause of psychiatric disorders such as ASD, schizophrenia or ADHD, which are tightly linked to the GABAergic system, and males are more likely to develop these pathologies. Preterm infants are 4 times more likely to develop ASD (Joseph et al., 2017). The possibility has been raised that this sex bias is an intrinsic female protective mechanism. This hypothesizes that multiple genetic factors contribute to the liability for developing ASD, and a higher threshold of genetic liability is required to result in female pathological presentation (Jacquemont et al., 2014). The “extreme male brain theory” is another prominent hypothesis to explain this gender bias, which suggests that fetal testosterone exposure may underlie gender differences in autistics traits (Baron-Cohen, 2002).

Since most of the perturbations observed in male preterm infants were related to the astrocytic regulation of GABAergic signaling, additional astrocytic transcripts related to either metabolism and morphogenesis or GABAergic system were analyzed. GABA-associated astrocytic genes were solely downregulated in males, oppositely to their normal developmental trajectory. For instance, glutamine and glutamate transporter transcripts were downregulated in preterm male infants. These neurotransmitter metabolisms are tied to the metabolism of GABA and their dysregulation has often been reported in psychiatric disorders (O’Donovan, Sullivan, & McCullumsmith, 2017).

In human, astrocytes are born during the second part of gestation (deAzevedo et al., 2003) leaving these cells highly susceptible to perinatal insults. Their proliferation occurs in tandem with birth and refinement of synapses within individual circuits, making astrocytes critical regulators of brain development (Perez-Catalan, Doe, & Ackerman, 2021). Since preterm male infants maintain the same number of astrocytes than their term counterpart, it seems that prematurity mainly affected the function of these cells through their maturation rather than their proliferation or survival. In accordance, preclinical studies highlighted the absence of reactive astrogliosis in the immature white matter following chronic hypoxia, with decreased glutamate transporter expression suggesting a more immature phenotype of astrocytes (Back et al., 2006; Raymond, Li, Mangin, Huntsman, & Gallo, 2011). However, the number of BA10-GABA-producing cells was decreased in preterm male infants, which is consistent with our former study (Lacaille et al., 2019) as well as other studies of preterm birth brain (Robinson et al., 2006) (Panda et al., 2018), and psychiatric disorders (Hashimoto et al., 2008) (Mellios et al., 2009) (Hashemi, Ariza, Rogers, Noctor, & Martínez-Cerdeño, 2017). Pharmacological effects of neuroprotective agents such as the GABA agonist and anti-inflammatory conpound allopregnanolone are being investigated in the context of preterm birth and show promising results(Vacher et al., 2021). New cellular models have been developped and will give the opportunity to investigate the environmental and genetic factors underlying injury in the developing human brain(Paşca et al., 2019). Even though prematurity is a significant contributor to psychiatric disorders, many preterm infants survive neurologically intact implying than most of the time, the preterm male infant GABA system could undergo a coordinated developmental delay, potentially conserving a stable excitatory-inhibitory balance.

## Contribution to the field

Extremely preterm birth carries a significantly increased risk of neurodevelopmental disorders, including autism and schizophrenia. A common feature of these disorders is an alteration of the GABAergic and glutamatergic neurotransmitter system in the prefrontal cortex, a region involved in working memory and social cognition. Here, we show that the maturation of the GABAergic system is more dynamic in male compared to in female term prefrontal cortex, potentially rendering males more susceptible to perinatal neurological insults. Astrocytic GABAergic system maturation was particularly delayed in male preterm prefrontal cortex. Defining a composite maturation index allowed meaningful temporal interpretation of cellular changes across variable human pathological samples. Understanding preterm prefrontal cortex alterations can provide insights into the perinatal environment’s impact on psychiatric disorders.

## Author contributions

H.L., C.-M.V. and A.A.P. conceived the project and designed the experiments. H.L. performed the experiments. H.L. and C.-MV. analyzed and interpreted the results. H.L., C.-M.V. and A.A.P. wrote, reviewed and revised the manuscript. All authors reviewed and revised the manuscript.

## Acknowledgements

We thank Dr. D. Bakalar, Dr J. O’reilly and J. Salzbank for insightful comments on the manuscript, and Janine Burgess and Sacha Parchment for expert assistance. We thank Dr. V. Gallo and Dr. E. Passegue for hosting us in their respective facilities. We also thank the contribution of families to the NIH NeuroBioBank, as well as J. Cottrell (NIH NeuroBioBank), who assisted us throughout the process of human tissue selection.

